# Genome-wide alternative splicing profiling in the fungal plant pathogen *Sclerotinia sclerotiorum* during the colonization of diverse host families

**DOI:** 10.1101/2020.05.13.094565

**Authors:** Heba M. M. Ibrahim, Stefan Kusch, Marie Didelon, Sylvain Raffaele

**Affiliations:** LIPM, Université de Toulouse, INRAE, CNRS, 31326 Castanet-Tolosan, France; Genetics Department, Faculty of Agriculture, Cairo University, 12613, Giza, Egypt; Plant Health and Protection, Division of Plant Biotechnics, Department of Biosystems (BIOSYST), Faculty of Bioscience Engineering, KU Leuven, Belgium; Unit of Plant Molecular Cell Biology, Institute for Biology I, RWTH Aachen University, 52056, Aachen, Germany

**Keywords:** *Sclerotinia sclerotiorum*, alternative splicing, host adaptation, computational analysis, isoforms, RNA sequencing (RNA-seq)

## Abstract

*Sclerotinia sclerotiorum* is a notorious generalist plant pathogen that threatens more than 600 host plants including wild and cultivated species. The molecular bases underlying the broad compatibility of *S. sclerotiorum* with its hosts is not fully elucidated. In contrast to higher plants and animals, alternative splicing (AS) is not well studied in plant pathogenic fungi. AS is a common regulated cellular process that increases cell protein and RNA diversity. In this study, we annotated spliceosome genes in the genome of *S. sclerotiorum* and characterized their expression *in vitro* and during the colonization of six host species. Several spliceosome genes were differentially expressed *in planta*, suggesting that AS was altered during infection. Using stringent parameters, we identified 1,487 *S. sclerotiorum* genes differentially expressed *in planta* and exhibiting alternative transcripts. The most common AS events during the colonization of all plants were retained introns and alternative 3′ receiver site. We identified *S. sclerotiorum* genes expressed *in planta* for which (i) the relative accumulation of alternative transcripts varies according to the host being colonized and (ii) alternative transcripts harbor distinct protein domains. This notably included 42 genes encoding predicted secreted proteins showing high confidence AS events. This study indicates that AS events are taking place in the plant pathogenic fungus *S. sclerotiorum* during the colonization of host plants and could generate functional diversity in the repertoire of proteins secreted by *S. sclerotiorum* during infection.

## Introduction

*Sclerotinia sclerotiorum* is a plant parasitic fungus that causes the white mold disease. It is known for its aggressive necrotrophic life style, which means that the fungus actively kills the plant host cells and thrives by feeding on the dead plant material, and for exhibiting a broad host range. *S. sclerotiorum* can infect more than 600 host plants including economically important species, such as tomato (*Solanum lycopersicum*), sunflower (*Helianthus annuus*), common bean (*Phaseolus vulgaris*), and beetroot (*Beta vulgaris*) (Boland and Hall, 1994; Naito and Sugimoto, 2011; Peltier et al., 2012).

Studies on broad host range plant pathogens largely focused on the function of virulence-related proteins and the transcriptional control of infection, but the molecular bases underlying the infection of diverse host plants are still not fully understood (Liang & Rollins, 2018). *S. sclerotiorum* synthesizes and secretes oxalic acid in order to establish successful colonization of host plants (Liang & Rollins, 2018). Further, *S. sclerotiorum* employs large numbers of cellulases, peptidases and toxins that assist in the necrotrophic infection process (Friesen *et al.*, 2008; Derbyshire *et al.*, 2017). Many plant pathogens employ secreted proteins functioning specifically on certain host genotypes to facilitate infection (Friesen *et al.*, 2008; Rodriguez-Moreno *et al.*, 2018), suggesting that the ability to infect very diverse host species would associate with expanded repertoires of secreted proteins. However, the repertoire of secreted protein coding genes in *S. sclerotiorum* is within the average for Ascomycete fungal pathogens (Derbyshire *et al.*, 2017). Instead of an expanded secretome, *S. sclerotiorum* exhibits codon usage optimization for secreted proteins, highlighting that a remarkably efficient protein translation system of virulence factors may support plant infection processes in *S. sclerotiorum* (Badet *et al.*, 2017). In addition, *S. sclerotiorum* hyphae organize in cooperating units, sharing the metabolic cost of virulence and growth during the colonization of resistant plants (Peyraud *et al.*, 2019). Here, we investigated the extent to which posttranscriptional regulation could generate diversity in virulence factor candidates produced by *S. sclerotiorum* during the colonization of plants from diverse botanical families.

Alternative splicing (AS) is a process in eukaryotic cells that increases the cellular capacity to shape their transcriptome diversity and proteome complexity. Splicing is an important mechanism that regulates the maturation of the precursor messenger RNAs (pre-mRNA) by subjecting it to the removal of non-coding sequences (introns). AS occurs in many eukaryotes under certain conditions resulting in multiple isoforms of transcripts that retain specific intronic sequences or lack specific exonic sequences. The transcripts with retained introns (RI) then have a prolonged lifetime compared to the completely mature mRNA transcript (Braunschweig *et al.*, 2014; Schmitz *et al.*, 2017; Naro *et al.*, 2017).

The efficiency and accuracy of the splicing mechanisms play a critical role in gene transcription and subsequent protein function. Imprecise splicing may result in abnormal and non-functional transcripts that may lead to the production of defective proteins, thus disturbing cellular processes. Previous studies showed that inaccurate splicing may cause diseases in humans and increases plant sensitivity to abiotic or biotic stresses (Cui *et al.*, 2014). In line with this, the importance of AS in plant immunity against pathogen attacks is well established (Rigo *et al.*, 2019). AS regulation and the factors that control it, the prediction of their *cis*-regulatory sequences and *trans*-acting elements have been intensively studied in plants and in animals (Blanco & Bernabeu, 2011; Eckardt, 2013; Zhang *et al.*, 2017), while only few reports are available from fungal phytopathogens. Therefore, the extent to which AS is regulated and functional during host colonization in fungal phytopathogens remains elusive.

Recently, (Jin *et al.*, 2017) found that transcripts of the plant fungal pathogen *Verticillium dahliae* undergo splicing of retained introns producing different isoforms of transcripts. These isoforms have predicted roles in controlling many conserved biological functions, such as ATP synthesis and signal transduction. The involvement and regulation of the retained intron isoforms and splicing during the infection of host plants are still unexplored. Moreover, AS is detected during *V. dahliae* microsclerotia development (Xiong *et al.*, 2014). Interestingly, 90% of the detected alternative transcripts exhibit retained introns. However, there is no further evidence to support the contribution of AS in microsclerotia development. In the same fashion, alternative transcripts are annotated in the genomes of the plant-pathogenic fungi *Colletotrichum graminicola* and *Fusarium graminearum* (Zhao *et al.*, 2013; Schliebner *et al.*, 2014).

Alternative splicing is pivotal in regulating gene expression and in diversification of the protein repertoire in the plant-pathogenic oomycete *Pseudoperonospora cubensis* during pathogen development and transition from sporangia to zoospores (Burkhardt *et al.*, 2015). In this study 4,205 out of 17,558 genes with *ca* 10,000 potential AS events were identified, of which *ca* 83% had evidence of retained introns. Interestingly, no exon skipping events were detected. Intriguingly, two genes encoding putative secreted RXLR and QXLR effectors showed evidence for retained intron specifically at the sporangia stage, while the spliced version was abundant during the host-associated stage. The retained intron may therefore regulate gene expression instead of affecting the function of the protein. Similarly, alternative splicing of the genes encoding glyceraldehyde-3-phosphate dehydrogenase (GAPDH) and 3-phosphoglycerate kinase (PGK) modulates their localization in the smut fungus *Ustilago maydis*. In particular, alternative splicing gives rise to GAPDH carrying a peroxisome targeting signal. Importantly, *U. maydis* mutants lacking the specific isoforms with peroxisomal localization have reduced virulence (Freitag *et al.*, 2012). These examples highlight the crucial role of AS in the pathogenicity of plant-pathogenic fungi.

A predicted splicing factor 8 corresponding to the U5-associated component Prp8 (GenBank accession number SS1G_03208) was reported recently from *S. sclerotiorum* (McLoughlin *et al.*, 2018). This prompted us to test for AS in *S. sclerotiorum* during the infection of diverse host plants. To this end, we exploited RNA-seq data of *S. sclerotiorum* infecting host plants from six botanical families, i.e. *Arabidopsis thaliana* (Brassicales), tomato (Solanales), sunflower (Asterales), beetroot (Caryophyllales), castor bean (*Ricinus communis*, Malphigiales), and common bean (Fabales), in addition to the RNA-seq of *S. sclerotiorum* cultivated *in vitro* as control (Peyraud *et al.*, 2019; Sucher *et al.*, 2020). We found that *S. sclerotiorum* has a functional splicing machinery and that at least 4% of the *S. sclerotiorum* secretome undergo alternative splicing regulation resulting in multiple differentially expressed isoforms that may have modified or altered functions. Some of the novel transcripts exhibit different predicted function or localization. Based on our analysis, we suggest that AS has the potential to give rise to transcriptional flexibility, thus contributing to the broad host spectrum of the plant pathogenic fungus *S. sclerotiorum*.

## Results

### *S. sclerotiorum* spliceosome is differentially regulated during host colonization

To study alternative splicing in the fungal plant pathogen *S. sclerotiorum, we* first searched the predicted proteome of *S. sclerotiorum* for components associated with splicing (spliceosome) using BLASTP and UniProtKB. We identified all the main components encompassing the entire pre-mRNA splicing cycle, i.e. U1/U2/U4/U5/U6-associated components, PRP19/NTC complex proteins, the proteins catalyzing the splicing of the intron (exon junction complex; EJC), the mRNA export complex TREX, and the mRNA and intron release components PRP43 and PRP22 (**Figure 1**).

**Figure 1.**
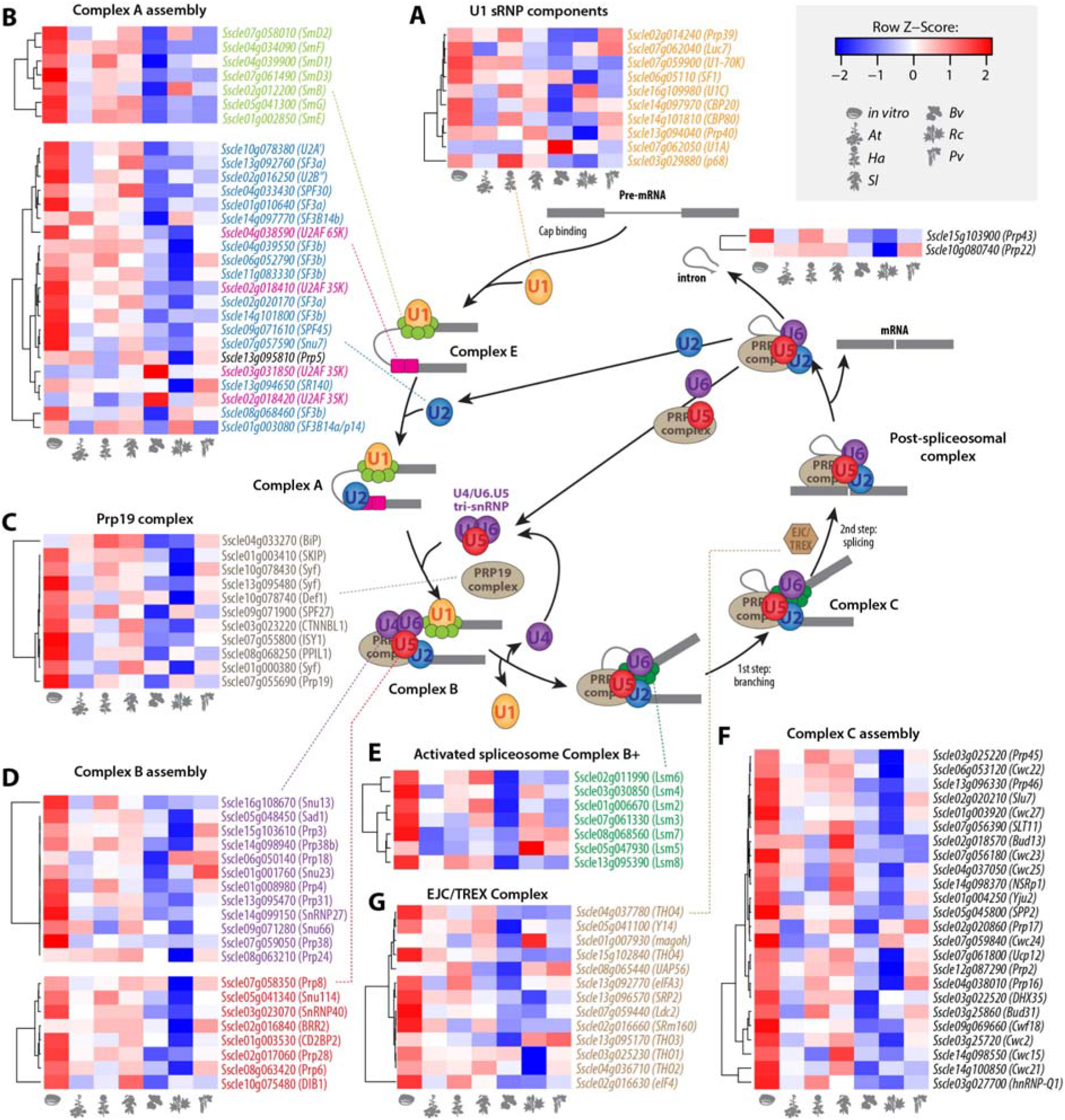
Identification of *S. sclerotiorum* spliceosome components and their transcriptional regulation during the infection of plants from six botanical families. Diagrammatic representation of mRNA splicing process featuring *S. sclerotiorum* genes involved in each step. A hypothetical pre-mRNA molecule is depicted with two exons shown as dark grey boxes and an intron shown as dark grey line. Circles, rounded rectangles and hexagon show protein complexes. The relative gene expression for 116 spliceosome genes at the edge of *S. sclerotiorum* mycelium during infection of plants from six species and *in vitro* is shown as heatmaps. Pre-mRNA splicing involves multiple spliceosomal complexes. First, complex E is established by binding of U1 snRNP (A) to small nuclear ribonucleoprotein-associated proteins (Sm), U2 associated factors (U2AF) and splicing factors (SF), leading to the recruitment of U2 and the formation of Complex A **(B).** The PRP19C/Prp19 complex/NTC/Nineteen complex **(C)** stabilizes the U4/U5/U6 tri-snRNP spliceosomal complex leading to Complex B assembly (D). The U1/U4 snRNPs are released to form the activated spliceosome complex B+ (E) triggering branching, intron excision, conformational rearrangements into complex C (F) and ligation of the proximal and distal exons. EJC/TREX is recruited to spliced mRNAs to mediate export to the cytoplasm **(G).** *At, Arabidopsis thaliana; Bv, Beta vulgaris; Ha, Helianthus annuus; Pv, Phaseolus vulgaris; Rc, Ricinus communis; Sl, Solanum lycopersicum*.

We documented the transcriptional regulation of *S. sclerotiorum* spliceosome components during plant infection by exploiting RNA-seq reads of *S. sclerotiorum* 1980 cultivated *in vitro* on PDA medium (Peyraud *et al.*, 2019) and during the infection of host plants from six botanical families (Sucher *et al.*, 2020): *A. thaliana* (*At*), tomato (*Solanum lycopersium, Sl*), sunflower (*Helianthus annuus, Ha*), common bean (*Phaseolus vulgaris, Pv*), castor bean (*Ricinus communis, Rc*), and beetroot (*Beta vulgaris, Bv*) (**Figure 1**). We found 116 proteins likely associated with (alternative) splicing, all but one of which were expressed at >10 FPKM across all conditions (**Figure 1; Supplementary Table 1**). *Sscle02g018420*, encoding a U2AF, was not expressed at detectable levels (FPKM < 1). By performing BLASTP searches we identified 81 of these 116 spliceosome-associated genes to be conserved in related *Ascomycetes*, such as *Botrytis* species (**Supplementary Table 2**). Interestingly, many components exhibited strongest expression *in vitro*, but appeared to be down-regulated on some or all of the hosts. Eighty of the 116 genes were significantly down-regulated (p<0.01) on at least one host plant, and one gene (*Sscle03g031850*, encoding a U2AF) was up-regulated on all hosts except sunflower (**Supplementary Table 3**). For example, 63 components were down-regulated on *B. vulgaris*, while the U2AF-encoding gene *Sscle03g031850* displayed 4.7-fold up-regulation during infection of *B. vulgaris* (**Figure 1**). Overall, 81 of the 116 components appeared to be differentially modulated dependent on the host plant, suggesting host plant-specific regulation of the spliceosome in *S. sclerotiorum*.

### Alternatively-spliced genes are differentially expressed during host infection

To search for alternative splicing events in the *S. sclerotiorum* transcriptome *in planta* and to reduce false discovery rate due to pipeline-dependent bias, we applied a stringent strategy based on two pipelines employing either transcriptome alignment or *de novo* transcriptome assembly (**Figure 2A**). Transcriptome alignment is a robust and effective method of characterizing transcripts that are mapped to a provided reference transcriptome (including isoforms with skipped exons) while *de novo* transcriptome assembly mainly focuses on recovering transcripts with segments of the genome that are missing from the transcriptome alignment method, including retained introns (Martin & Wang, 2011).

**Figure 2.**
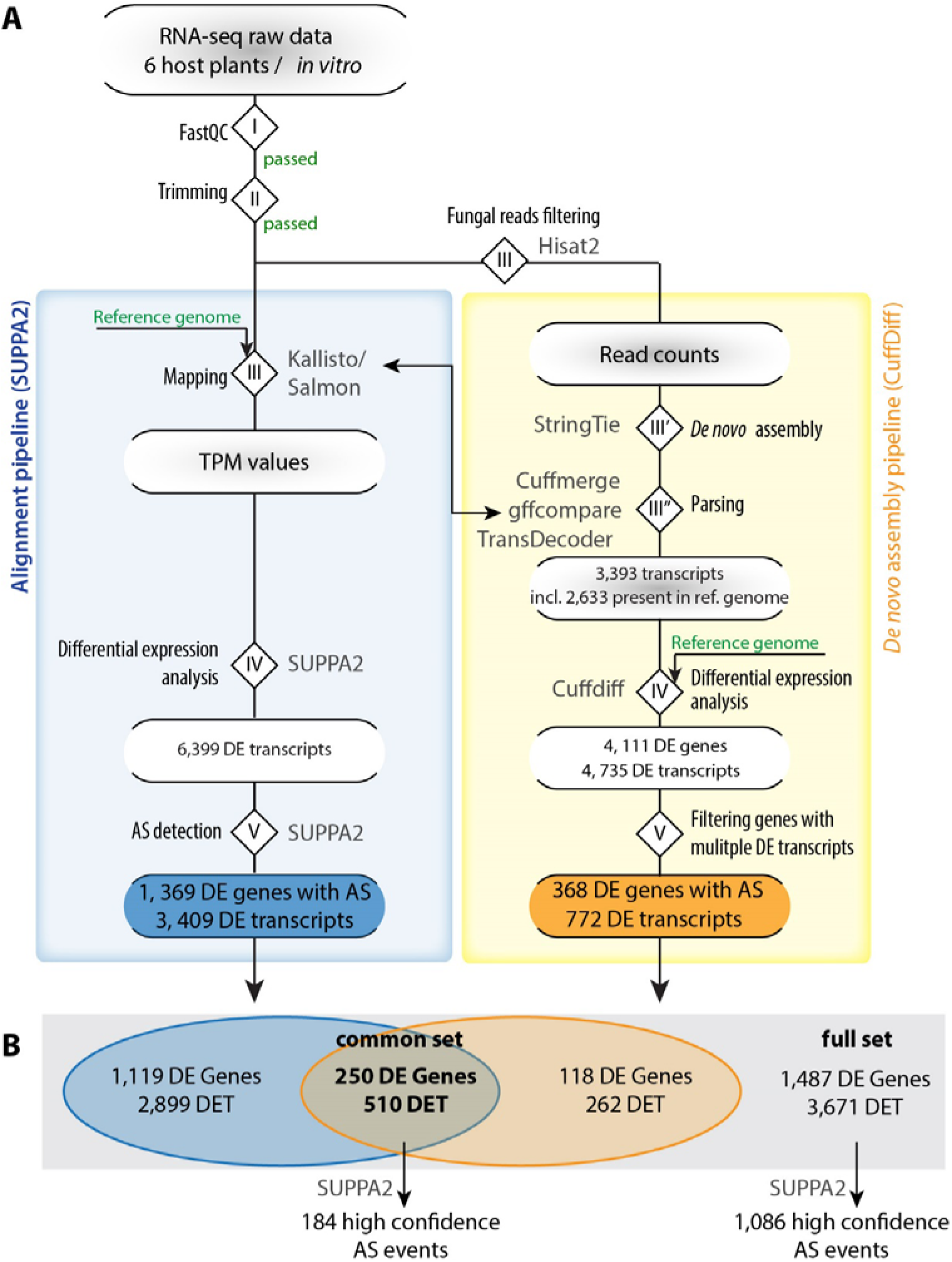
Pipeline for genome-wide detection of alternative splicing in *S. sclerotiorum*. **(A)** Raw RNA-seq data was first inspected with FastQC (I) and quality-trimmed using Trimmomatic (II). We then applied two pipelines for detection and analysis of novel transcripts, the *de novo* assembly pipeline (yellow box) and the transcriptome alignment pipeline (blue box). For detection of novel transcripts, we mapped reads with HISAT (III) to the *S. sclerotiorum* reference genome; this data was used in a modified StringTie *de novo* assembly (III’). Using Cuffmerge, gffcompare and Transdecoder, we identified novel transcripts compared with the reference gene annotation (III”) and generated a new reference annotation. Using the new annotation and the reference genome, we performed differential expression analysis (IV) and filtered the differentially expressed (DE) genes for those encoding at least two DE transcripts (DET; V). In the transcriptome alignment pipeline, mapping was done with Kallisto or Salmon (III), DE analysis and alternative splicing detection with SUPPA2 (IV and V). (B) A venn diagram summarizing the results from DE analysis in (A) for both pipelines; numbers are given only for genes encoding multiple transcripts. AS, alternative splicing; DE, differentially expressed; DET, differentially expressed transcript; incl., including; ref., reference; TPM, transcripts per million.

In the transcriptome alignment pipeline (**Figure 2A**), the trimmed reads (steps I and II) were aligned to the *S. sclerotiorum* 1980 reference genome (Derbyshire *et al.*, 2017) (step III). Expression of transcripts (transcripts per million, TPM) was determined in Salmon with the QUASI mapping algorithm (Patro *et al.*, 2017) and Kallisto (Bray *et al.*, 2016). Next, we used SUPPA2 to identify differentially expressed (DE) transcripts (step IV), with a cut-off TPM >30 and p-value <0.05 (Trincado *et al.*, 2018). We found 6,399 DE transcripts in total with this approach among all samples (lesion edge on six plant species) and compared to the control (edge *S. sclerotiorum* cultivated *in vitro* on PDA medium). Then, SUPPA2 was applied to DE genes to identify the different alternative splicing (AS) events and to measure the percent spliced in index (PSI; ψ), which represent the ratio between reads excluding or including exons (step V). These PSI values indicate the inclusion of sequences into transcripts (Wang *et al.*, 2008; Alamancos *et al.*, 2015) using the normalized transcript abundance values (TPM) of the isoforms from Salmon. The differential splicing analysis of the events (dpsi values) at p<0.05 identified 1,369 DE genes with significant splicing events producing 3,409 DE transcripts (**Figure 2A, B**).

In the *de novo* assembly pipeline, transcripts were assembled from fungal reads using StringTie. To identify fungal reads in our samples, the trimmed reads (step I and II) were aligned to the *S. sclerotiorum* 1980 reference genome (Derbyshire *et al.*, 2017) using HISAT2 (Kim *et al.*, 2015), yielding between 10,258,270 and 26,314,353 mapped reads per sample (**Supplementary Table 4**) (step III). *S. sclerotiorum* reads were then used for *de novo* transcriptome assembly in a modified Tuxedo differential expression analysis pipeline (Trapnell *et al.*, 2010, 2012). Since StringTie was proven to be a more accurate and improved transcript assembler and quantifier (Pertea *et al.*, 2015, 2016), we used StringTie instead of cufflinks for the *de novo* assembly step (step III’ and III”). This resulted in 3,393 transcripts, including 2,633 transcripts from genes present in the reference transcriptome of *S. sclerotiorum* isolate 1980 (Derbyshire *et al.*, 2017), 410 gene fusions and 337 novel genes encoding 350 transcripts. Differential expression analysis on the complete transcriptome including both reference and novel transcripts with cuffdiff (step IV) identified 4,111 DE genes accounting for 4,735 DE transcripts on any of the six host species compared to the control. Out of those, there were 368 genes that encoded several DE transcripts each, producing 772 transcripts in total. These represent candidate genes harboring alternative splicing *in planta* (step V).

Finally, we compared transcripts identified with the two pipelines and found a total number of 3,671 transcripts differentially expressed *in planta* in total, originating from 1,487 genes (‘full set’ of candidates). Among those, the two pipelines identified a common set of 250 genes of *S. sclerotiorum* encoding more than one transcript and expressed differentially *in planta* (‘common set’ of candidates, **Figure 2B** and **Supplementary Figure 1**). This common set of genes produced 510 transcripts differentially expressed *in planta*. To homogenize alternative splicing event predictions on these genes, we re-run SUPPA2 to calculate PSI values for genes from the common and full sets of candidates. This identified 1,086 high confidence AS events in the full set of genes and 184 high confidence AS events in the common set of genes.

### The alternative splicing landscape in *S. sclerotiorum* during host colonization

To document the effect of AS on *S. sclerotiorum* genes differentially expressed *in planta, we* performed AS events detection with SUPPA2 on genes induced on each plant, and classified AS by type of event on each plant. The number of AS events varied 2.63-fold according to host, ranging from 158 AS events in *A. thaliana* to 415 AS events in *B. vulgaris*, reaching a total 1,086 distinct AS events for the six plant species (**Figure 3A**). The distribution of AS event type did not differ significantly during colonization of the six different host plants (**Figure 3A**). Retained intron was the major type of AS event detected in *S. sclerotiorum* during host colonization (RI; 39.8±1.1%), followed by alternative 3′ receiver site (A3; 30.3±1.2%), alternative 5’ donor site (A5;15.0±06%), skipped exon (SE; 4.4%±1.3%) and alternative first exon (AF; 1.1±0.2%).

**Figure 3.**
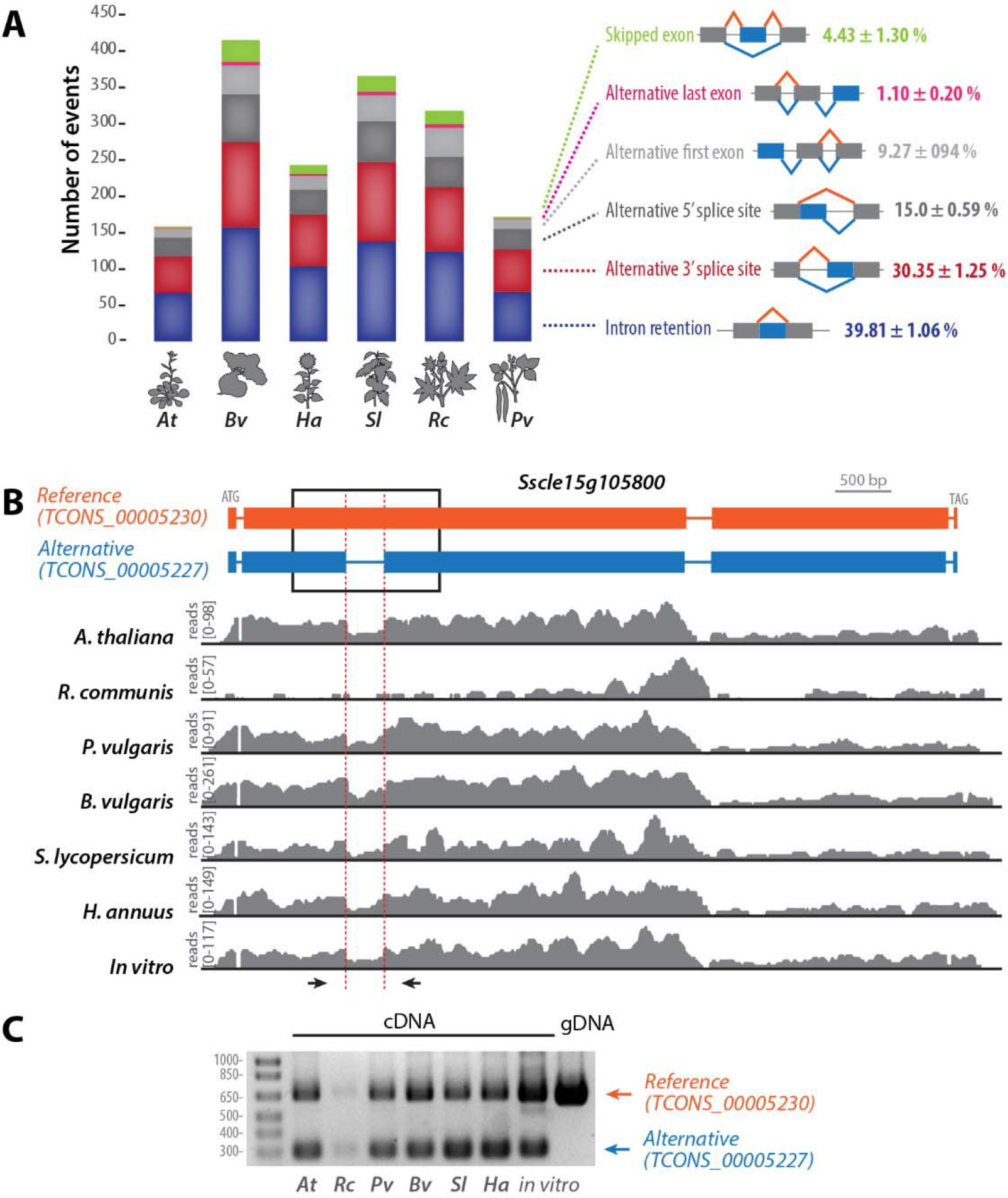
Intron retention is the major type of alternative splicing (AS) event in *S. sclerotiorum* during the colonization of plants from diverse botanical families. **(A)** Distribution of high confidence AS events identified by SUPPA2 according to type of event and host plant infected by *S. sclerotiorum*. In diagrams depicting the different type of events, orange lines show intron splicing pattern in the reference transcript, blue lines show intron splicing pattern in the alternative transcript, blue boxes show alternatively spliced exons, gray boxes show invariant exons. Percentages indicate the relative proportion of one AS event type relative to all AS events identified during infection of a given plant species. **(B)** Example of an intron retention event in the reference transcript of *Sscle15g105800*. In the transcripts diagram, exons are shown as boxes, introns as lines. Read mappings are shown in grey for one RNA-seq sample of each treatment. **(C)** RT-PCR analysis of *Sscle15g105800* transcripts produced during the colonization of six plant species and *in vitro*. The position of oligonucleotide primers used for RT-PCR is shown as arrows in (B). *At, Arabidopsis thaliana; Bv, Beta vulgaris; Ha, Helianthus annuus; Pv, Phaseolus vulgaris; Rc, Ricinus communis; Sl, Solanum lycopersicum*.

In all hosts, the most frequent AS event was intron retention, of which the gene *Sscle15g105800* is one example. This gene belongs to the full set of AS gene candidates and encodes a 2,043 amino-acids long predicted protein of unknown function conserved in *B. cinerea*. StringTie identified for this gene a transcript TCONS_00005227 with five exons and four introns (**Figure 3B**). The reference transcript TCONS_00005230 harbors four exons, including a 3,958 bp exon 2 corresponding to the fusion between TCONS_0005227 exon 2, retained intron 2 (351 bp), and exon 3. Reads aligned to TCONS_0005227 intron 2 were detected in all RNA-seq samples, and were particularly abundant during infection of *A. thaliana, B. vulgaris* and *P. vulgaris. Sscle15g105800* was weakly expressed on *R. communis* with few reads aligned to TCONS_0005227 intron 2 (**Figure 3B**). To confirm alternative splicing of TCONS_0005227 intron 2, we performed RT-PCR with primers spanning this intron on genomic DNA and on cDNAs produced from *S. sclerotiorum* grown *in vitro* (PDA medium) and infected plants (**Figure 3C**). Amplicons from the reference transcript TCONS_00005230 were detected on all cDNAs, albeit only weakly on cDNAs produced from infected *R. communis*. An alternative transcript was detected on all cDNAs the size of which corresponds to TCONS_00005227 retained intron 2.

### Alternative splicing is host-regulated in *S. sclerotiorum*

To document the extent to which host plant species associated with alternative splicing events in *S. sclerotiorum, we* performed hierarchical clustering and principal component analysis (PCA) for *S. sclerotiorum* alternatively spliced transcript accumulation in six host species (**Figure 4A**). The distribution of the plant variable according to the two principal components displayed host-specific clustering, in which AS transcripts produced on each host could be clearly separated, except for AS transcripts produced on *A. thaliana* and *S. lycopersicum*. This analysis suggested that the relative accumulation of alternative transcripts produced by a given gene could vary according to the host being colonized.

**Figure 4.**
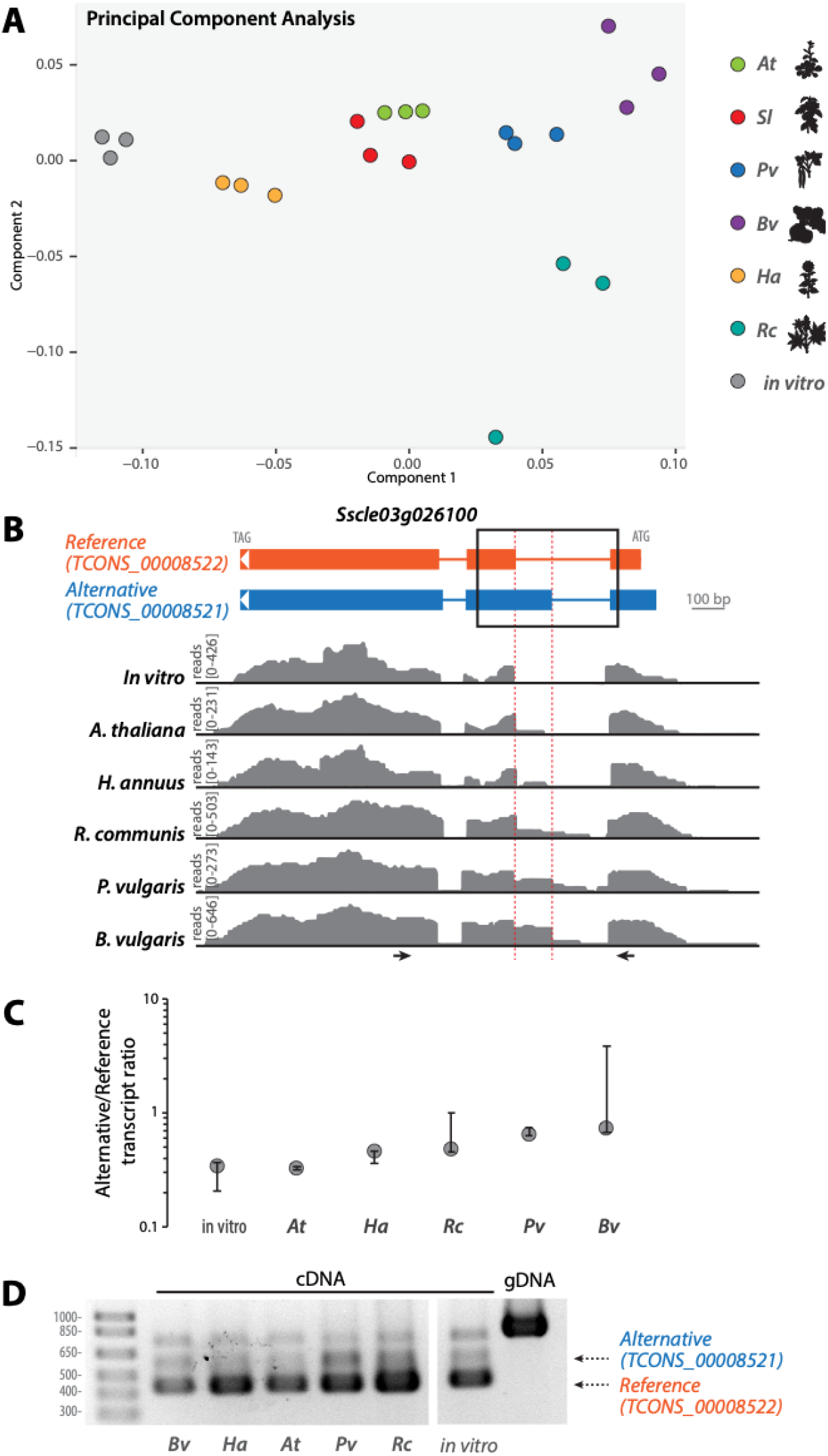
Alternative 3’ receiver splice site variation according to host in *Sscle03g026100*. **(A)** Principal component analysis map of the sample variable for the accumulation of reference and alternative transcripts produced by 250 high confidence *S. sclerotiorum* genes showing alternative splicing. Sample types are color-coded according to infected host plant species. (B) Example of alternative 3’ receiver splice site in the reference transcript of *Sscle03g026100*. In the transcript diagrams, exons are shown as boxes, introns as lines. Read mappings are shown in grey for one RNA-seq sample of each treatment. (C) Ratio between the abundance of alternative over reference transcript for *Sscle03g026100* determined by Cuffdiff. Error bars show 90% confidence interval. (D) RT-PCR analysis of *Sscle03g026100* transcripts produced during the colonization of five plant species and *in vitro*. The position of oligonucleotide primers used for RT-PCR is shown as arrows in (B). *At, Arabidopsis thaliana; Bv, Beta vulgaris; Ha, Helianthus annuus; Pv, Phaseolus vulgaris; Rc, Ricinus communis; Sl, Solanum lycopersicum*.

We tested whether this was the case for the gene *Sscle03g026100*, encoding a predicted phosphoenolpyruvate kinase-like protein. The *Sscle03g026100* locus harbored RNA-seq reads that aligned in the 3’ region of intron 1, indicative of alternative 3’ receiver sites in exon 2 of the reference transcripts (*TCONS_00008522*, **Figure 4B**). This splicing event is predicted to cause an extension of the alternatively spliced exon in transcript variant *TCONS_00008521*. Thanks to its N-terminal extension, the protein isoform TCONS_0008522 but not TCONS_00008521 is recognized as a member of the PIRSF034452 family (TIM-barrel signal transduction protein). According to Cuffdiff transcript quantification, the ratio between alternative and reference transcript varied from 0.32 in *A. thaliana* to 0.73 in *B. vulgaris* (**Figure 4C**). To confirm alternative splicing of *Sscle03g026100* transcript, we performed RT-PCR with primers spanning the variant exon 2 on RNAs collected from five host species (**Figure 4D**). We retrieved a 418 bp amplicon corresponding to the reference transcript (*TCONS_00008522*), a 535 bp amplicon corresponding to the *TCONS_00008521* alternative transcript, as well as a third ~750 bp amplicon. In agreement with the RNA-seq read coverage, bands corresponding to the alternative transcript *TCONS_0008521* were much weaker than bands corresponding to the reference transcript *TCONS_0008522* in *A. thaliana, H. annuus* and *in vitro*.

### Alternative splicing is predicted to generate protein isoforms with modified functions

To study the functional consequences of AS in *S. sclerotiorum*, we first analyzed gene ontology (GO) terms enriched in our list of high confidence DE genes with AS. GO enrichment was determined using BiNGO, a tool package within the complex network visualizing platform “Cytoscape” (Shannon *et al.*, 2003; Maere *et al.*, 2005) (**Supplementary Table 5**). The most significantly enriched terms included “oxidoreductase activity” and “carbohydrate metabolic process”, suggesting that genes involved in the degradation of carbohydrates and organic molecules were subject to alternative splicing during infection of host plants. According to BLASTP searches (E value <1E-25) 175 of the 250 genes in our common set of AS candidates are conserved in related *Ascomycetes*, including *Botrytis* species (**Supplementary Table 6**).

Second, to test if AS could alter the domain content of protein isoforms in our full set of AS candidates, we assigned PFAM domains to all isoforms and identified AS events leading to a change in PFAM domain content. In total, 158 genes expressed alternative transcripts with changes in PFAM annotation profiles. Of these, 53 isoforms exhibited loss of PFAM domains, 85 isoforms displayed gain of PFAM domains, and 20 isoforms show more complex changes of PFAM profiles (**Supplementary File 1**). Only 8 of the 158 genes with alterations in their PFAM annotation profiles were from the 250 common AS genes (**Table 1**). Four isoforms gained PFAM domains, e.g. the putative cutinase Sscle11g080920 where the alternative isoform TCONS_00002255 gained two PFAM domains, ETS_PEA3_N (PF04621.12) and CBM_1 (PF00734.17).

**Table 1:**
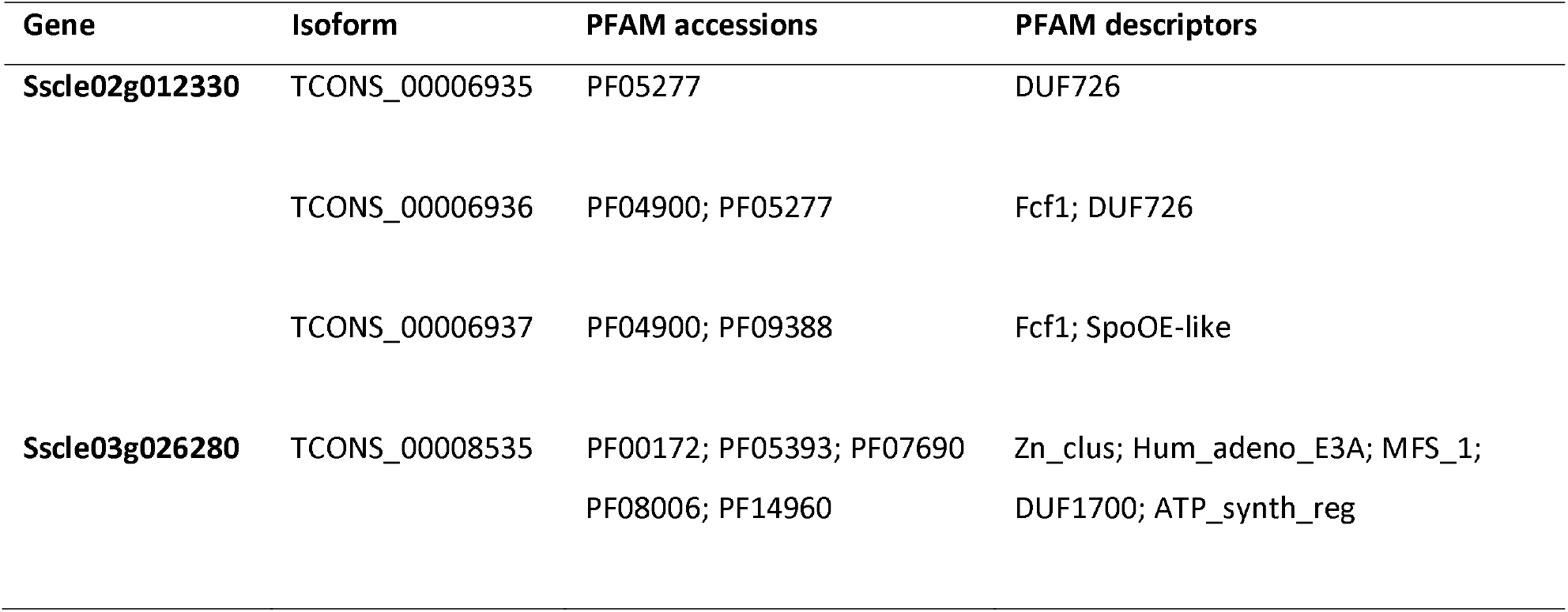
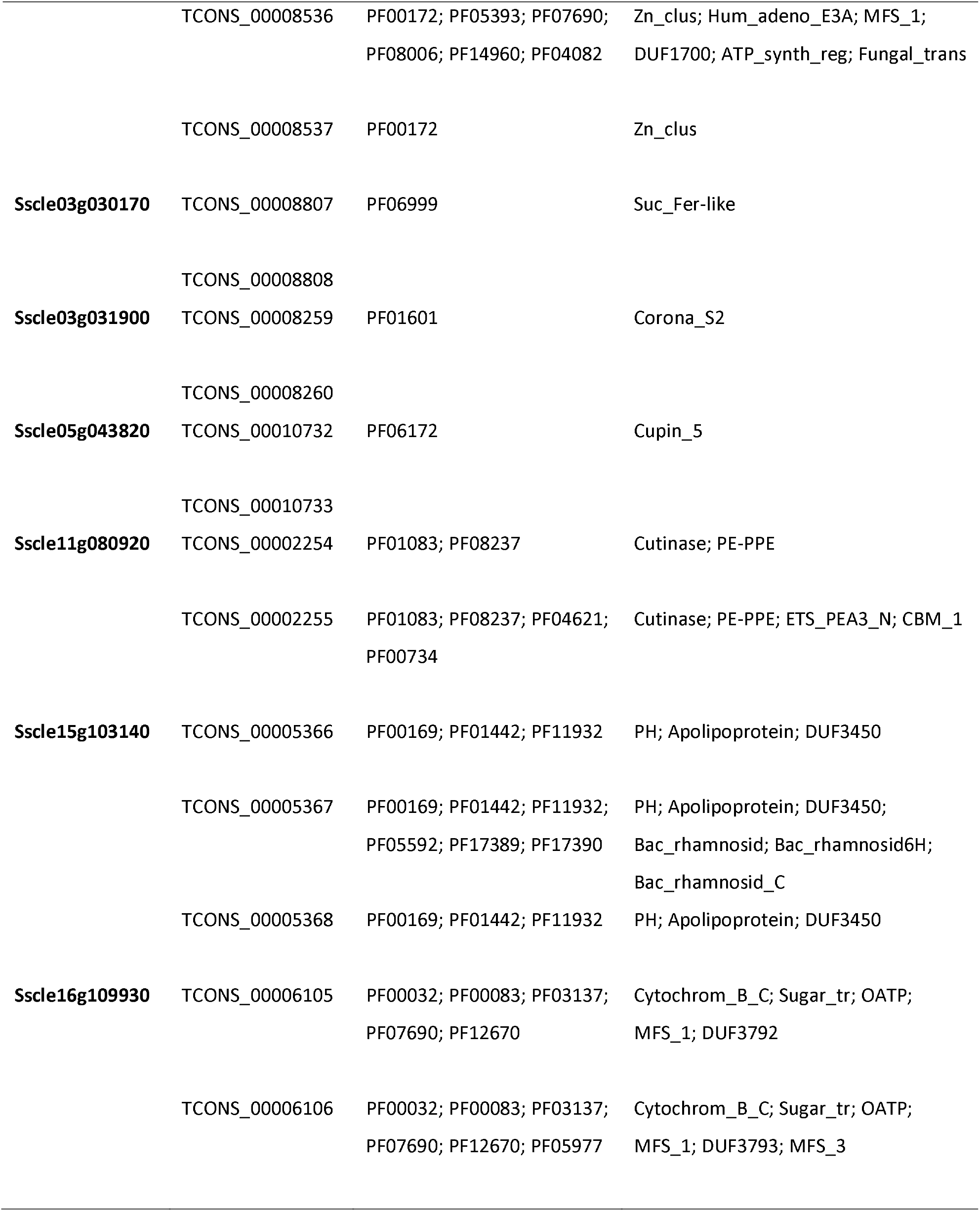
Changes in PFAM profiles of differentially expressed and alternatively spliced genes of *S. sclerotiorum*.

Furthermore, we explored the isoforms from alternatively spliced genes for signal peptides for secretion (**Figure 5A**). Of the 250 genes in our common set of AS candidates, 42 are predicted to encode a secreted protein, corresponding to 4% of the *S. sclerotiorum* secretome (Juan et al., 2019; **Supplementary Figure 2**). Among those, five genes (*Sscle02g014060, Sscle07g057820, Sscle09g070580, Sscle12g091110*, and *Sscle15g103140*) showed possible gains of secretion peptide by AS and two cases of loss of secretion peptides in alternative isoforms (*Sscle10g075480* and *Sscle15g102380*). In the set of 3,393 AS candidates detected in total, we found 26 possible gains of secretion peptides and 16 losses of secretion peptides in alternative isoforms.

**Figure 5.**
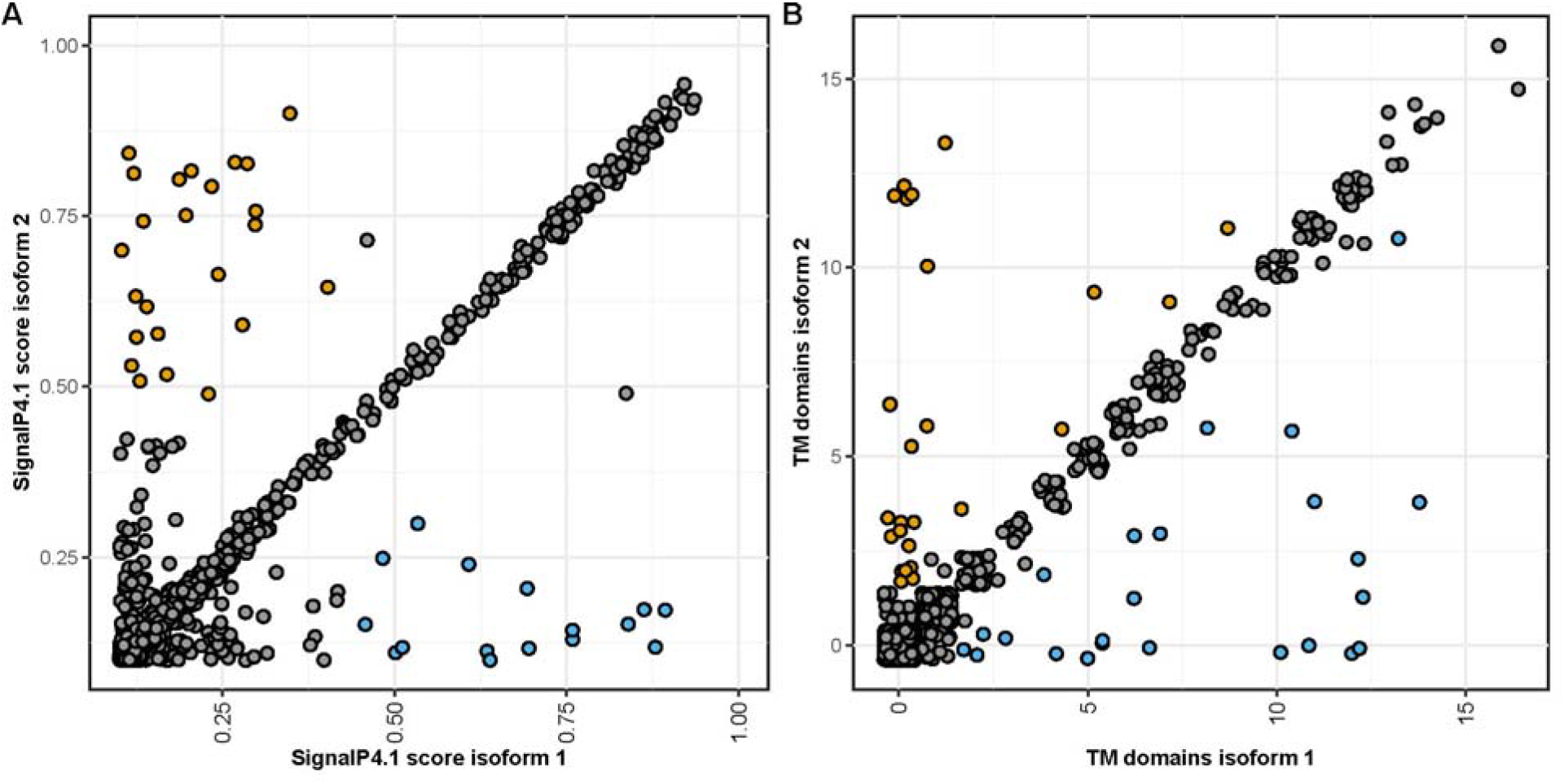
Alternative splicing modifies secretion potential of proteins in *S. sclerotiorum*. **(A)** We determined the likelihood of presence of an N-terminal secretion signal using SignalP4.1 in the protein encoded by the reference transcript (isoform 1) and the alternative transcripts (isoform 2). The scatterplot shows the SignalP scores of both isoforms for all alternatively spliced genes of *S. sclerotiorum*, where a score of 0.45 is the threshold for a putative secretion peptide. Orange data points indicate novel isoforms that may have gained a secretion peptide, blue data points indicate loss of the secretion peptide. (B) We predicted the number of transmembrane domains for reference (isoform 1) and novel isoforms (isoform 2) using TMHMM. Orange data points indicate isoforms that gained transmembrane domains, blue data points indicate novel isoforms that may have lost one or more transmembrane domains.

Similarly, alternative splicing caused the gain and loss of transmembrane domains in novel isoforms. In total, 20 novel isoforms exhibited gain of one and 25 isoforms gained more than one predicted transmembrane domains. 32 of these did not harbor a putative transmembrane domain in the reference isoform, which suggests re-localization to the plasma membrane or an intracellular membrane. *Vice versa, we* observed the loss of one transmembrane domain in 30 isoforms and of more than one in 24 isoforms, including 32 isoforms that lost all transmembrane domains, suggesting subcellular re-localization of the respective novel isoform. Two genes who gained (*Sscle05g040780* and *Sscle16g109930*) and five genes that lost (*Sscle02g014060, Sscle02g019060, Sscle03g031900, Sscle05g043820, Sscle08g066940*) transmembrane domains are found in the 250 alternatively spliced and differentially expressed genes. Intriguingly, the novel isoform of Sscle02g014060 is predicted to be secreted as well as to have lost its transmembrane domain.

### Alternative splicing is predicted to modify the activity of *S. sclerotiorum* secreted proteins

Of the 250 genes with evidence for AS, 42 are predicted to encode a secreted protein, corresponding to 4% of the *S. sclerotiorum* secretome (Juan et al., 2019; **Supplementary Figure 2**). For example, the alternatively spliced gene *Sscle11g080920* was predicted to encode for two secreted protein isoforms derived from the reference transcript TCONS_00002255 and the alternative transcript TCONS_00002254, exhibiting an alternative 5’ donor splice site in exon 3 (**Figure 6A**). The relative accumulation of these two isoforms varied according to the plant being colonized. The reference transcript was expressed more strongly than the alternative transcript in *S. sclerotiorum* infecting *A. thaliana* (**Figure 6B**). By contrast, we measured higher accumulation of the alternative than the reference transcript in *S. sclerotiorum* infecting *P. vulgaris*. To confirm alternative splicing of the *Sscle11g080920* transcript, we performed RT-PCR with primers spanning the variant exon 3 on RNA collected from *A. thaliana* and *P. vulgaris* (**Figure 6C**). As expected, this identified a 618 bp amplicon corresponding to the reference transcript and a 430 bp amplicon corresponding to the alternative transcript during the colonization of *A. thaliana* and *P. vulgaris*. In this assay, the alternative transcript accumulated more than the reference transcript during the infection of *P. vulgaris*, and to a lesser extent during the colonization of *A. thaliana*.

**Figure 6.**
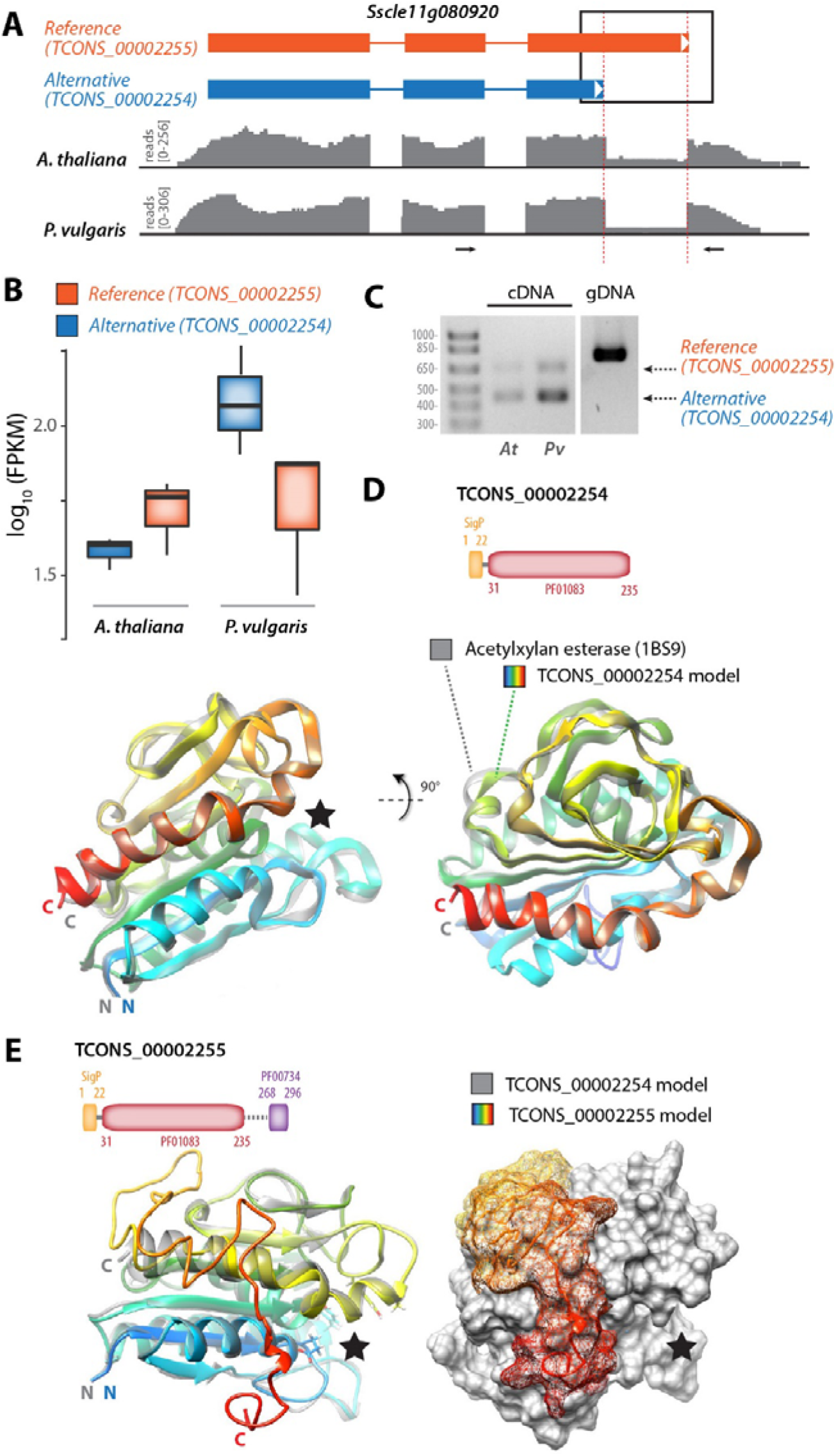
Alternative splicing generates structural diversity in the predicted secreted protein Sscle11g080920. **(A)** Consequences of alternative 5’ donor splice site in exon 3 of the reference transcript of *Sscle11g080920*. In the transcript diagrams, exons are shown as boxes, introns as lines. Read mappings are shown in grey for one RNA-seq sample collected on infected *A. thaliana* and infected *P. vulgaris*. **(B)** Relative accumulation of the reference and alternative transcripts produced by *Sscle11g080920* on infected *A. thaliana* and infected *P. vulgaris*. Boxplots show the expression of the transcripts TCONS_00002254 and TCONS00002255 from RNA-seq of *S. sclerotiorum* infecting *A. thaliana* and *P. vulgaris* in log10(FPKM). Boxplots show the median of the data points, whiskers are at 1.5x interquartile range of the highest/lowest value. **(C)** RT-PCR analysis of *Sscle11g080920* transcripts produced during the colonization of *A. thaliana* (*At*) and *P. vulgaris* (*Pv*). The position of oligonucleotide primers used for RT-PCR is shown as arrows in (A). **(D)** Diagram of the domain structure for the protein produced by TCONS_00002254 alternative transcript, and ribbon model of TCONS_00002254 predicted protein structure (rainbow colors). TCONSS_00002254 protein model is shown superimposed with its closest structural analog AXEII acetylxylan esterase (grey). The black star indicates the position of the active site in AXEII. **(E)** Diagram of the domain structure for the protein produced by TCONS_00002255 reference transcript, and ribbon model of TCONS_00002255 predicted protein structure (rainbow colors). TCONSS_00002255 protein model is shown superimposed with TCONS_00002254 model (grey). The black star indicates the position of the active site deduced from the analysis shown in (D).

To gain insights into the functional consequence of AS in *Sscle11g080920, we* analyzed PFAM domains and performed structure modelling for the proteins encoded by the reference transcript TCONS_00002255 and the alternative transcript TCONS_00002254. The alternative transcript TCONS_00002254 encoded a 235 amino acid protein featuring a secretion signal and a cutinase (PF01083) domain (Figure 6D). Homology modeling and fold recognition in I-TASSER identified the acetylxylan esterase AXEII from *Penicillium purpureogenum* (PDB identifier 1BS9) as the closest structural analog to TCONS_00002254 (RMSD 0.31Å). AXEII is a close structural analog of *Fusarium solani* cutinase, an esterase that hydrolyzes cutin in the plant’s cuticle (Ghosh *et al.*, 2001). The reference transcript TCONS_00002255 encoded a 296 amino-acid protein featuring a secretion signal, a cutinase domain and a short fungal cellulose binding domain (PF00734) (Figure 6E). Its closest structural analog identified by I-TASSER was model 1G66 of AXEII. The superimposition of TCONS_00002254 and TCONS_00002255 protein models revealed that the C-terminal extension in TCONS_00002255 corresponds to a surface-exposed unstructured loop reaching the neighborhood of the catalytic site cleft (Figure 6E). This additional exposed loop could modify protein-protein interactions in TCONS_00002255 or modify access to its catalytic site. These results suggest that alternative splicing is a mechanism to generate functional diversity in the repertoire of proteins secreted by *S. sclerotiorum* during the colonization of host plants.

## Discussion

There are several approaches to study alternative splicing from RNA-seq data (Thakur *et al.*, 2019), such as analyzing splice junctions (Hu *et al.*, 2013) or exonic regions (Anders *et al.*, 2012), which largely rely on mapping strategies only. The pipeline we used in this study combines two fundamentally different strategies (*de novo* assembly-based and reference mapping-based) to detect true novel splicing events and reduce algorithm bias. This approach however does not completely exclude false positive or false negative alternative splicing events, and also does not allow to distinguish between an alternative splicing event and correction of an incorrect reference gene model. Manual inspection or curation of gene models, as for example performed in *F. graminearum* (Zhao *et al.*, 2013), is required to distinguish between these possibilities. We have inspected AS predictions and experimentally validated alternative transcripts for a small subset of the AS events predicted here, supporting the accuracy of our analysis pipeline. Nevertheless, further efforts will be needed to improve the gene annotations of *S. sclerotiorum*, confirm alternative transcripts, and identify further alternatively spliced transcripts missed by our pipeline at the genome scale.

### A number of *S. sclerotiorum* genes are spliced alternatively on different hosts

Colonization of a host plant by a pathogen requires global changes in gene expression of the pathogen and secretion of effector proteins and enzymes (van der Does & Rep, 2017). Alternative splicing is a regulatory mechanism affecting the activity of a majority of genes in plant and animal cells at the post-transcriptional level. Whether AS contributes to the regulation of virulence in plant-pathogenic fungi remains elusive. In this study, we present a comparative genome-wide survey of AS in the plant-pathogenic fungus *S. sclerotiorum* during the infection of six different host plants compared to growth *in vitro* as a control. Using stringent criteria for the detection of alternatively spliced isoforms, and considering genes identified consistently with our two pipelines, we found 250 genes that expressed more than one isoform (**Figure 2**). These represent about 2.3% of the genome, which is consistent with estimates for the AS rate in the closely related fungi species *Botrytis cinerea* (Grützmann *et al.*, 2014). In *Fusarium graminearum*, alternative splicing represents 1.7% of the total number of genes (Zhao *et al.*, 2013), while in *Colletotrichum graminicola* only 0.57% of the total number of the predicted genes appear to exhibit alternative splicing (Schliebner *et al.*, 2014). Yet, this percentage is strikingly low compared to the AS rate of the intron or multiexon-containing genes in plants such as *Arabidopsis thaliana* or mammals such as *Homo sapiens*, which are reported to be 42% and 95%, respectively (Pan *et al.*, 2008; Filichkin *et al.*, 2010). AS is not well characterized in plant-pathogenic fungi and needs to be investigated in more detail (Grützmann *et al.*, 2014). A previous study reported evidence for AS in the plant-pathogenic oomycete *Pseudoperonospora cubensis* that causes downy mildew in the *Cucurbitaceae* family (Burkhardt *et al.*, 2015). In this work, 24% of the expressed genes showed novel isoforms with new AS events over the course of infection of cucumber at 1-8 days after infection. Moreover, recently Jin et al. (2017) found that the transcripts of two different isolates of the plant fungal pathogen *Verticillium dahliae* undergo splicing of retained introns producing different isoforms of transcripts that differ between the two isolates during the fungal development. These isoforms have predicted roles in controlling many conserved biological functions, such as ATP synthesis and signal transduction.

In our analysis, most of the alternative splicing events were retained introns (RI; 39.8%), which is consistent with previous studies were intron retention showed higher preference in the newly identified isoforms (Grützmann *et al.*, 2014). On the other hand, skipped exons represent a small frequency in our analysis (4.4%) but could be considered higher than usual compared to other fungi such as *Verticillium dahliae* (2-fold higher; 2.2%). Interestingly, SE is the most common AS event in mammals (Sammeth *et al.*, 2008).

### Do these AS variants contribute to virulence on the respective hosts?

Alternative splicing is a natural phenomenon in eukaryotes that is genetically tightly regulated, and proper spliceosome activity ensures adequate splicing (Chen *et al.*, 2012). The operating mechanisms of splicing regulation and the extent to which components of the splicing machinery regulate splice site decisions remain poorly understood however (Saltzman *et al.*, 2011). The spliceosome activity is modulated by *cis*- and *trans*-acting regulatory factors. The *trans*-acting elements include the SR (serine/arginine-rich) and hnRNP (heterogeneous ribonucleoprotein) families (Chen & Manley, 2009; Nilsen & Graveley, 2010) and generally regulate AS by enhancing or inhibiting the assembly of the spliceosome at adjacent splice sites after perceiving *cis*-acting elements in exon or intron regions of pre-mRNAs.

Since spliceosome components strictly regulate splicing, any changes in spliceosome component abundance may result in inaccurate splicing and/or generation of alternative transcripts in accordance with the environmental condition that causes the changes. Although tightly regulated, AS is influenced by external stimuli in eukaryotes such as biotic and abiotic stresses. For instance, the LSM2–8 complex and SmE, which are regulatory components of the spliceosome, differentially modulate adaptation in response to abiotic stress conditions in Arabidopsis (Carrasco-López *et al.*, 2017; Huertas *et al.*, 2019). Similarly, U1A is essential in adapting Arabidopsis plants to salt stress. Mutation in AtU1A renders Arabidopsis plants hypersensitive to salt stress and results in ROS accumulation (Gu *et al.*, 2018).

In our analysis we found that many of the spliceosome components are down-regulated in *S. sclerotiorum* during infection, in particular on hosts where we detected a high number of AS events. For instance, during infection of *B. vulgaris S. sclerotiorum* exhibited the highest abundance of alternative transcripts and showed down-regulation of the majority of the spliceosome components (**Fig. 1** and **Fig. 3**). This suggests that AS in *S. sclerotiorum* could result from the down-regulation of spliceosome genes and raises the question of whether host plant defenses actively interfere with the regulation of *S. sclerotiorum* spliceosomal machinery to trigger the observed down-regulation. A related goal for future research will be determining whether alternative splicing confers fitness benefits to *S. sclerotiorum* during host colonization. Dedicated functional analyses will be required to clarify the role of AS in *S. sclerotiorum* adaptation to host.

In some cases, more than one isoform is present in a host plant. The reason could be that the new isoforms have new functions that assist the establishment of pathogenesis, while the dominant isoform(s) has/have substantial biological functions that are needed for *Sclerotinia* under any condition. The different isoforms produced by AS in *S. sclerotiorum* during infection of the different host plants could be a way to increase pathogen virulence. For instance, the alternatively spliced gene *Sscle03g026100* encodes a putative phosphonopyruvate hydrolase. Phosphonopyruvate hydrolases hydrolyze phosphonopyruvate (P-pyr) into Pyruvate and Phosphate (Liu *et al.*, 2004). In plants, phosphonopyruvate plays an important intermediate role in the formation of organophosphonates, which function as antibiotics and play a role in pathogenesis or signaling. Therefore, the fungus may use these two different isoforms to detoxify one of the plant defense molecules in order to facilitate the infection process. In a previous study, *Ochrobactrumanthropi* and *Achromobacter* bacterial strains were found to degrade the organophosphates from surrounding environments and to use the degraded product as a source of carbon and nitrogen (Ermakova *et al.*, 2017). Interestingly, the newly identified isoform of Sscle03g026100 (TCONS_00008521) showed the highest expression during infection of beans (*B. vulgaris*). Since *B. vulgaris* is well known for their production of antifungal secondary metabolites such as C-glycosyl flavonoids and betalains (Citores *et al.*, 2016; Ninfali *et al.*, 2017), this suggests that the new isoform may be required for *S. sclerotiorum* to overcome the plant resistance by degrading some of these metabolites. In addition, alternative splicing *of Sscle11g080920*, predicted to encode a secreted cutinase, could exhibit specificity for differently branched cellulose molecules. Taken together, our study revealed that *S. sclerotiorum* uses alternative splicing that gives rise to functionally divergent proteins. We further show that a number of these isoforms have differential expression on diverse host plants.

## Material and Methods

### Plant inoculations and RNA sequencing

Raw RNA-seq data used in this work is available from the NCBI Gene Expression Omnibus under accession numbers GSE106811, GSE116194 and GSE138039. Samples and RNAs were prepared as described in (Sucher *et al.*, 2020). Briefly, the edge of 25 mm-wide developed necrotic lesions were isolated with a scalpel blade and immediately frozen in liquid nitrogen. Samples were harvested before lesions reached 25 mm width, at 24 hours (*H. annuus*), 47 to 50 hours (*A. thaliana, P. vulgaris, R. communis* and *S. lycopersicum*) or 72 hours post inoculation (*B.vulgaris*). Material obtained from leaves of three plants were pooled together for each sample, all samples were collected in triplicates. RNA extractions were performed using NucleoSpin RNA extraction kits (Macherey-Nagel, Düren, Germany) following the manufacturer’s instructions. RNA sequencing was outsourced to Fasteris SA (Switzerland) to produce Illumina single end reads (*A. thaliana, S. lycopersicum, in vitro* control) or paired reads (other infected plants) using a HiSeq 2500 instrument.

### Quantification of isoform and transcript abundance

Quality control for the RNA-seq data was performed using FastQC (Babraham Bioinformatics). The quality-checked data were processed for trimming with the Java-based tool Trimmomatic-0.36 (Bolger et al., 2014). Transcript abundances were quantified using a set of tools as follows: In the alignment pipeline, reads were first mapped to the *S. sclerotiorum* reference genome (Derbyshire et al. 2017) using HISAT2 (Kim *et al.*, 2015). Annotation of reference genes and transcripts were provided in the input. The aligned reads were assembled and the transcripts were quantified in each sample using StringTie (Pertea *et al.*, 2015, 2016). The assemblies produced by StringTie were merged with the reference annotation file in one GTF file to incorporate the novel isoforms with the original ones using cuffmerge (Goff *et al.*, 2019). The accuracy of the merged assembly was estimated by reciprocal comparison to the *S. sclerotiorum* reference annotation. In the *de novo* assembly pipeline, transcript abundances were quantified using gffcompare (Pertea *et al.*, 2016) and cuffcompare (Trapnell *et al.*, 2012). All the expressed transcripts including novel genes and alternatively spliced transcripts were merged into one annotation file using the Tuxedo pipeline merging tool, cuffmerge. The accuracy of the assembled annotation file was assessed by comparing it to the reference genome using gffcompare (Pertea *et al.*, 2016).

### Differential expression analysis of RNA-seq

The differential expression analysis of genes and isoforms were calculated using cuffdiff from the Tuxedo pipeline (Trapnell *et al.*, 2012). We then used CummeRbund to visualize the cuffdiff results of the genes whose expression were marked as significant and at log2 fold change of ±2 across all samples, leaving 4,111 genes that had differentially expressed isoforms (**Figure 2**). *quasi-mapping* was applied on the same RNA-seq data for expression quantification of transcripts using Kallisto (Bray *et al.*, 2016) and Salmon-0.7.0 (Patro *et al.*, 2017). Differential expression analysis of the quantified transcript and isoform abundance of the RNA-seq data resulting from StringTie and Salmon, were used in cuffdiff (Trapnell *et al.*, 2012) and SUPPA2 (Trincado *et al.*, 2018), respectively, according to the default parameters as referred to by the software manuals. The R Studio software packages CummeRbund (Goff *et al.*, 2019) was employed to determine the significant change in the transcript abundance across the different samples. All samples were compared with the PDA *in vitro* cultivation control. Default settings were used. Genes with an FDR-adjusted *P* (*q*) < 0.05 with a fold change of ±2 were considered differentially expressed.

### RNA-seq data visualization and transcripts annotation

The Integrative Genomics Viewer (IGV) (Robinson *et al.*, 2017) and Web Apollo annotator (Lee *et al.*, 2013) were used for visualizing the RNA-seq data. Heatmaps were generated with the heatmap.2 function of R (R Core Team 2018). Spliceosome genes were identified using several approaches. A first set of genes were identified based on map 03040 (Spliceosome) for *S. sclerotiorum* (organism code ‘ssl’) in the Kyoto Encyclopedia of Genes and Genomes (KEGG). The annotation of these genes was verified using BLASTP searches against *Saccharomyces cerevisiae* and *Homo sapiens* in the NCBI ReSeq database followed by searches in the UniprotKB database for detailed annotation. Second, we searched for all spliceosome components annotated in Ascomycete genomes in the UniprotKB database and identified their orthologs in *S. sclerotiorum* using BLASTP searches. Gene ontology classification database with the Blast2GO package was used to perform the functional clustering of the differentially expressed or spliced genes. The method was performed using Fisher’s exact test with robust false discovery rate (FDR) correction to obtain an adjusted P-value between certain tested gene groups and the whole annotation. SignalP4.1 (Nielsen, 2017) was used to predict N-terminal secretion signals of reference and novel isoforms. Transmembrane domains were predicted with TMHMM v2.0 (Krogh *et al.*, 2001).

### Reverse transcription PCR

RNAs were collected as for the RNAseq experiment. Reverse transcription was performed using 0.5 μL of SuperScriptII reverse transcriptase (Invitrogen), 1 μg of oligo(dT), 10 nmol of dNTP and 1 μg of total RNA in a 20 μL reaction. RNAs collected from 3 plants of each species were pooled together for cDNA synthesis. RT-PCR was performed using gene-specific primers (**Supplementary Table 7**) on an Eppendorf G-storm GS2 mastercycler with PCR conditions 4 min at 94°C followed by 32 cycles of 30 s at 94°C, 30 s at 55°C and 1 min at 72°C, followed by 10 min at 72°C.

### Protein 3D structure modeling and visualization

Protein structure models were determined using the I-TASSER online server (Yang *et al.*, 2015). Top protein models retrieved from I-TASSER searches were rendered using the UCSF Chimera 1.11.2 software. Models were superimposed using the MatchMaker function in Chimera, best-aligning pair of chains with the Needleman-Wunsch algorithm with BLOSUM-62 matrix and iterating by pruning atom pairs until no pair exceeds 2.0 angstrom.

## Conflict of Interest

The authors declare that the research was conducted in the absence of any commercial or financial relationships that could be construed as a potential conflict of interest.

## Author contributions

HMMI suggested the idea, designed the experiment, performed the bioinformatics analyses, and drafted the manuscript. SK contributed to the bioinformatics analyses and drafted the manuscript. MD performed the RT-PCR. SR monitored the RT-PCR experiment, contributed to bioinformatics analyses, revised the manuscript and provided feedback. All authors edited and proofread and approved the final version of this manuscript.

## Funding

This work was supported by a Starting grant from the European Research Council (ERC-StG-336808) to SR and the French Laboratory of Excellence project TULIP (ANR-10-LABX-41; ANR-11-IDEX-0002-02).

## Acknowledgements

This work is dedicated to the memory of Mohammed M. I. Al-Maraghy, who had recently passed away. This work was supported by the French and the Egyptian governments through a co-financed fellowship by the French Embassy in Egypt (Institute Français d’Egypte; IFE) and the Science and Technology Development Fund (STDF). We are grateful to the genotoul bioinformatics platform Toulouse Midi-Pyrenees (Bioinfo Genotoul, http://bioinfo.genotoul.fr/) for providing help and/or computing and/or storage resources.

## Data availability statement

All datasets generated for this study are included in the manuscript and the supplementary files. RNA-seq data is deposited at NCBI GEO under accessions GSE106811, GSE116194 and GSE138039.

## Notes

https://www.ncbi.nlm.nih.gov/bioproject/PRJNA418121

https://www.ncbi.nlm.nih.gov/bioproject/PRJNA477716

https://www.ncbi.nlm.nih.gov/bioproject/PRJNA574280

### Competing Interest Statement

The authors have declared no competing interest.

