## Supplementary figures and images for "Genome-wide alternative splicing profiling in the fungal plant pathogen *Sclerotinia sclerotiorum* during the colonization of diverse host families"

### Supplementary Figure 1

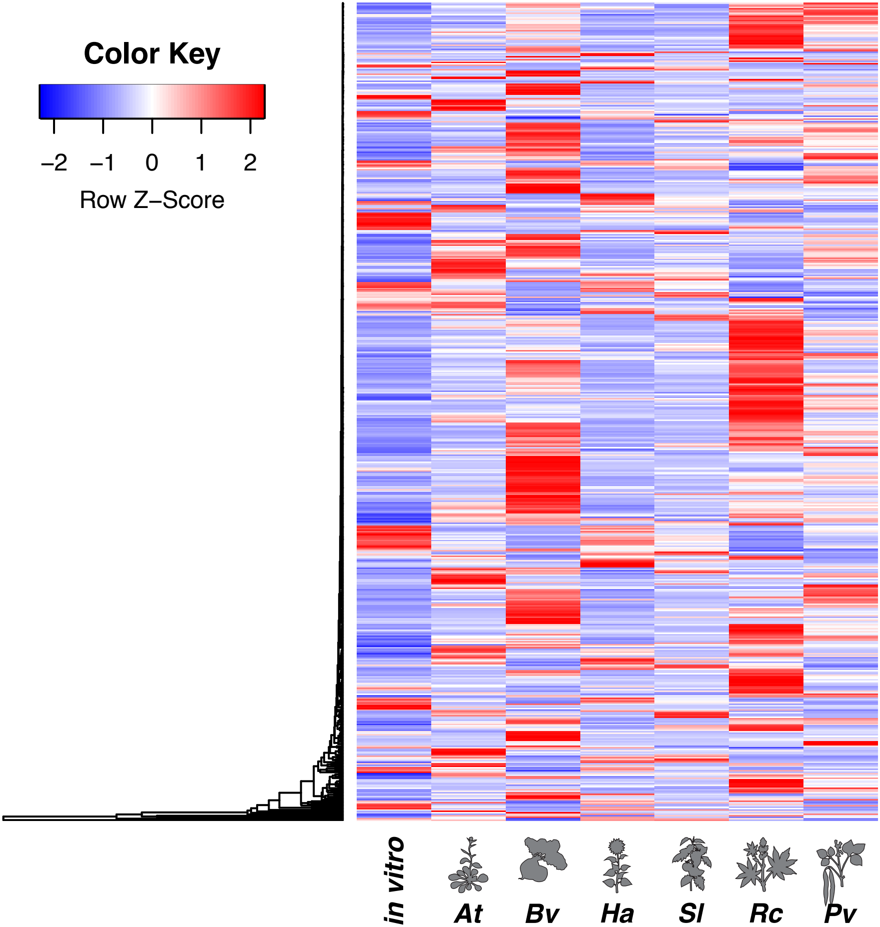

### Supplementary Figure 2

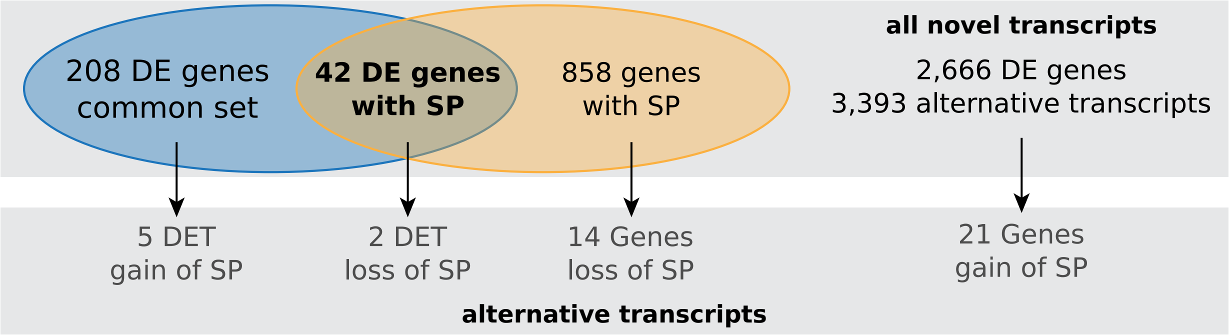
